# High-dimensional regression over disease subgroups

**DOI:** 10.1101/092825

**Authors:** Frank Dondelinger, Sach Mukherjee, The Alzheimer’s Disease Neuroimaging Initiative

## Abstract

We consider high-dimensional regression over subgroups of observations. Our work is motivated by biomedical problems, where disease subtypes, for example, may differ with respect to underlying regression models, but sample sizes at the subgroup-level may be limited. We focus on the case in which subgroup-specific models may be expected to be similar but not necessarily identical. Our approach is to treat subgroups as related problem instances and jointly estimate subgroup-specific regression coefficients. This is done in a penalized framework, combining an *ℓ*_1_ term with an additional term that penalizes differences between subgroup-specific coefficients. This gives solutions that are globally sparse but that allow information-sharing between the subgroups. We present algorithms for estimation and empirical results on simulated data and using Alzheimer’s disease, amyotrophic lateral sclerosis and cancer datasets. These examples demonstrate the gains our approach can offer in terms of prediction and the ability to estimate subgroup-specific sparsity patterns.

## 1 Introduction

High-dimensional regression has been well studied in the case where all samples can reasonably be expected to follow the same model. However, in several current and emerging applications, observations span multiple subgroups that may not be identical with respect to the underlying regression models. Examples abound, from disease subtypes in biomedicine to customer subsets in business applications. We are specifically motivated by biomedical problems, where sets of samples, such as disease subtypes, although related, may differ with respect to underlying biology and therefore have different relationships between covariates and a response of interest.

Thus, we focus on high-dimensional regression in group-structured settings. In particular, we consider linear regression in a commonly-encountered scenario in which the same set of *p* covariates or predictors is available in each of *K* subgroups. That is, we consider subgroup-specific linear regression problems indexed by *k*, each with subgroup-specific sample size *n_k_*, a response vector *y_k_* of length *n_k_*, a *n_k_* × *p* feature matrix *X_k_* and a *p*-vector *β_k_* of regression coefficients. The problem we address is estimating the regression coefficients *β*_1_…β_*К*_.

We propose an approach to jointly estimate the regression coefficients that induces global sparsity and encourages similarity between subgroup-specific coefficients. We consider the following penalized formulation and its variants

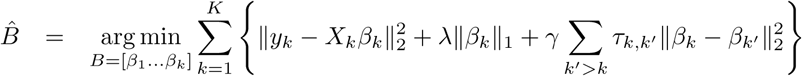

where, *B* = [*β*_1_…*β*_*К*_] is a *p* × *K* matrix that collects together all the regression coefficients, ∥ ˙ ∥_*q*_ denotes the *ℓ*_*q*_ norm of its argument and *λ*, *γ*, *τ* are tuning parameters. The last term is a fusion-type penalty between subgroups; note that the difference is taken between entire vectors of subgroup-specific coefficients. An *ℓ*_2_ fusion penalty is shown above, although other penalties may be used; in this manuscript, we also consider an *ℓ*_1_ variant. The parameters *τ*_*k*_,*k*′ allow for the possibility of controlling the extent to which similarity is encouraged for specific pairs of subgroups.

Our proposal shares similarities with both the group lasso [Yuan and Lin, 2006] and the fused lasso [Tibshirani et al., 2005] but differs from both in important ways. In contrast to the group lasso, we consider subgroups of samples or observations rather than groups of coefficients and in contrast to the fused lasso, we consider fusion of entire (subgroup-specific) coefficient vectors, rather than successive coefficients under a pre-defined ordering. Obozinski et al. [2010] showed how the group lasso could be used in subgroup-structured settings, essentially by considering the global problem and defining groups (in the group lasso sense) corresponding to the same covariate across all subgroups. This means that each covariate tends either to be included in all subgroup-specific models or none. In contrast, our approach allows subgroups to have different sparsity patterns, whilst pulling subgroup-specific coefficients together and inducing global sparsity. Our work is also similar in spirit to recent work concerning joint estimation of graphical models over multiple problem instances [Danaher et al., 2014, Oates et al., 2014, 2015].

We are motivated by emerging problems in biomedical research and specifically in personalized medicine. High-dimensional regression problems are now becoming common in this area, with several high-dimensional data types already in mainstream use. In the personalized medicine setting, samples usually correspond to individuals and the subgroups *k* to e.g. diseases or disease subtypes. It is increasingly clear that many disease subtypes differ in their biology [see e.g. Weinstein et al., 2013, Akbani et al., 2014], suggesting that relationships between covariates and responses of interest may differ between them. However, sample sizes tend to be limited, especially at the subgroup level, posing problems for the subgroup-wise strategy of solving each problem separately. On the other hand, pooling all the data together into a single regression problem may lead to severe mis-specification if the underlying subgroup-specific models do indeed differ.

These issues may lead to losses in terms of predictive ability and perhaps just as important in the ability to efficiently estimate subgroup-specific influences that may themselves be of interest. In contrast to simple pooling, our approach allows subgroups to have different sparsity patterns and regression coefficients, but in contrast to the subgroup-wise approach it takes advantage of similarities between subgroups.

We show empirical results in the context of two neurodegenerative diseases - Alzheimer’s disease and amyotrophic lateral sclerosis (ALS) - and cancer (see below for full details of the applications and data). The responses concern disease progression in Alzheimer’s and ALS and therapeutic response in cancer cell lines. In the Alzheimer’s and ALS examples, subgroups are based on clinical factors, while in the cancer data they are based on the tissue type of the cell lines.

Across the three examples, data types include genetic, clinical and transcriptomic variables. We find that our approach can improve performance relative to pooling or subgroup-wise analysis. Importantly, in cases where pooling or subgroup-wise analyses do well (perhaps reflecting a lack of subgroup structure or insufficient similarity respectively) our approach remains competitive. This gives assurance that penalization is indeed able to share information appropriately in real-world examples. We emphasize that the goal of the empirical analyses we present is not to give the best predictions possible in these applications, but rather to explore joint estimation in group-structured biomedical problems.

## 2 Methods

### 2.1 Notation

Each subgroup *k* ∈ {1… *K*} has the same set of *p* covariates, but subgroup specific sample size *n_k_*. Total sample size is 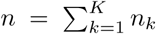 For subgroup *k*,*X_k_* is the *n_k_* × *p* feature matrix and *y_k_* the corresponding *n_k_* × 1 vector of observed responses. Subgroup-specific regression coefficients are *β_k_* ∈ ℝ^*p*^. Where convenient we collect all regression coefficients together in a *p* × *k* matrix *B* = [*β*_1_…*β_k_*]and accordingly we use *β_j_*,*_k_* to denote the coefficient for covariate *j* in subgroup *k*.

### 2.2 Model Formulation

We seek to jointly estimate the regression coefficients *B* = [*β*_1_ … *β_k_*] whilst ensuring global sparsity and encouraging agreement between subgroup-specific coefficients. We propose the criterion

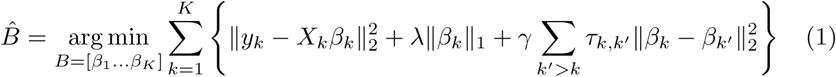

and a variant with an *ℓ*_1_ norm in the last term

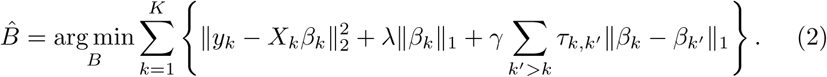

Here, *λ*, *γ*, *τ* are tuning parameters. The role of the last term is to encourage similarity between subgroup-specific regression coefficients. The special case *K* = 1 recovers the classical lasso (applied to all data pooled together). The tuning parameters *τ_k_*,_*k′*_ give the possibility of controlling the extent of fusion between specific subgroups. By default all *τ*’s are set to unity (“unweighted fusion”), but they can also be set to specific values as discussed below (“weighted fusion”). In the above formulation, we assume that *y_k_* and *X_k_* have been standardized (at the subgroup level) so that no intercept terms are required. Note that the regularization parameters *λ*, *γ* are the same across subgroups.

The difference between the two variants is that the first, *ℓ*_2_ fusion encourages similarity between subgroup-specific coefficients, while the second *ℓ*_1_ version allows for exact equality. The *ℓ*_2_ formulation has the computational advantage that the fusion part of the objective function becomes continuously differentiable, and the estimate of the objective function at each step can be obtained by soft-thresholding, analogously to coordinate descent for regular lasso problems. In the *ℓ*_1_ formulation on the other hand, the fusion constraint is only piece-wise continuously differentiable, leading to a more difficult optimisation problem (see below).

### 2.3 Comparison with group and fused lasso

Our formulation resembles the group lasso and fused lasso, but differs from both in important ways. The original group lasso [Yuan and Lin, 2006] was designed to consider groups of covariates within a single regression problem. Let *X* be the feature matrix and *y* the vector of responses in a standard regression problem. Letting *l ∊* {1… *L*} index groups of covariates, the group lasso criterion is

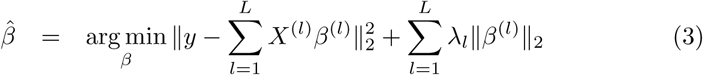

where *X^(ℓ)^* is the submatrix of *X* corresponding to the covariates in group *ℓ*,*β*^(*ℓ*)^ the corresponding regression coefficients and *λ*_l_’s tuning parameters. The penalty tends to include or exclude all members of a group from the model, i.e. all coefficients in a group may be set to zero giving groupwise sparsity.

In our setting, the subgroups are subsets of samples rather than covariates. Nevertheless, as shown in Obozinski et al. [2010], one could use a group lasso-like criterion for estimation in the multiple subgroup setting by forming groups *l* each comprising all the coefficients for a single covariate *j ∈* {*1*… *p*} across all *K* regression problems. This encourages covariates to either be included in all the subgroup-specific models or none.

The fused lasso [Tibshirani et al., 2005] is also aimed at a single regression problem, but assumes that the covariates can be ordered in such a way that successive coefficients may be expected to be similar. This leads to the following criterion

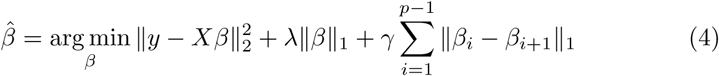

where *λ*,*γ*are tuning parameters and we have assumed that the covariates are in a suitable order. The final term encourages similarity between successive coefficients. Efficient solutions for various classes of this problem exist [e.g. Hoefling, 2010, Liu et al., 2010, Ye and Xie, 2011].

Our approach shares the use of a fusion-type penalty, but focuses on a different problem, namely that of jointly estimating regression coefficients across multiple, potentially non-identical, problems. Accordingly, our penalty encourages agreement between entire coefficient vectors from different subgroups and does not require any ordering of covariates.

### 2.4 Setting the tuning parameters *τ*

For weighted fusion, the parameters *τ_k_*,*_k_*′ could be set by cross-validation but this may be onerous in practice. As an alternative we consider setting *τ_k_*,*_k_*′ using a distance function *d*(*k*,*k*′) based on the covariates. The idea is to allow more fusion between subgroups that are similar with respect to *d*, while allowing the *τ_k_*,*_k′_* to be set in advance of estimation proper. However, this assumes that similarity in the covariates reasonably reflects similarity between the underlying regression coefficients, which may or may not be the case in specific applications.

We consider two variants. The first sets *d*(*k*,*k*′) = ∥*μ_k_* – *μ_k_*′∥2 where *μ_k_*,*μ_k_*′ are the sample means of the covariates in the subgroups *k*,*k*′respectively (we assume the data have been standardized). The second approach additionally takes the covariance structure into account by using the symmetrised Kullback Leibler (KL) divergence, i.e. 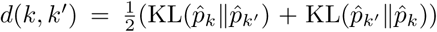, where 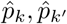 are estimated marginal distributions over the covariates in the subgroups *k*, *k*′ respectively and KL(*p*∥*q*) is the KL-divergence between distributions *p* and *q*. In practice, this requires simplifying distributional assumptions. Below we use multivariate Normal models for this purpose, with the graphical lasso [Friedman et al., 2008] used to estimate the Σ*_k_*’s. For both approaches, we set *τ_k_*,*_k_*′ = 1 — *d*(*k*,*k*′)/*d*_max_, with *d_max_* the largest distance between any pair of groups *k*,*k*′ (this scales *τ* to the unit interval).

### 2.5 Optimisation

We describe a coordinate descent approach for optimising equation (1). While it is possible to derive a block coordinate descent approach for equation (2) [e.g. following Friedman et al., 2007], this is generally inefficient for the highdimensional problems that we consider. Instead, we will describe an optimization procedure based on a proximal gradient approximation derived in Chen et al. [2010].

#### 2.5.1 Coordinate Descent for *ℓ*_2_ Fusion

The *ℓ*_2_ fusion penalty is continuously differentiable and we can obtain the optimal value for 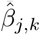 in equation (1) at each step by first calculating optimal values without the lasso penalty:

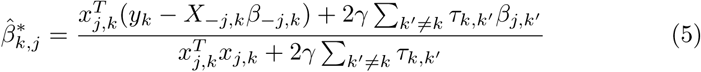

Then 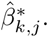 can be obtained by soft-thresholding on 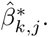). The procedure is summarized in Algorithm 1.

**Algorithm 1.**
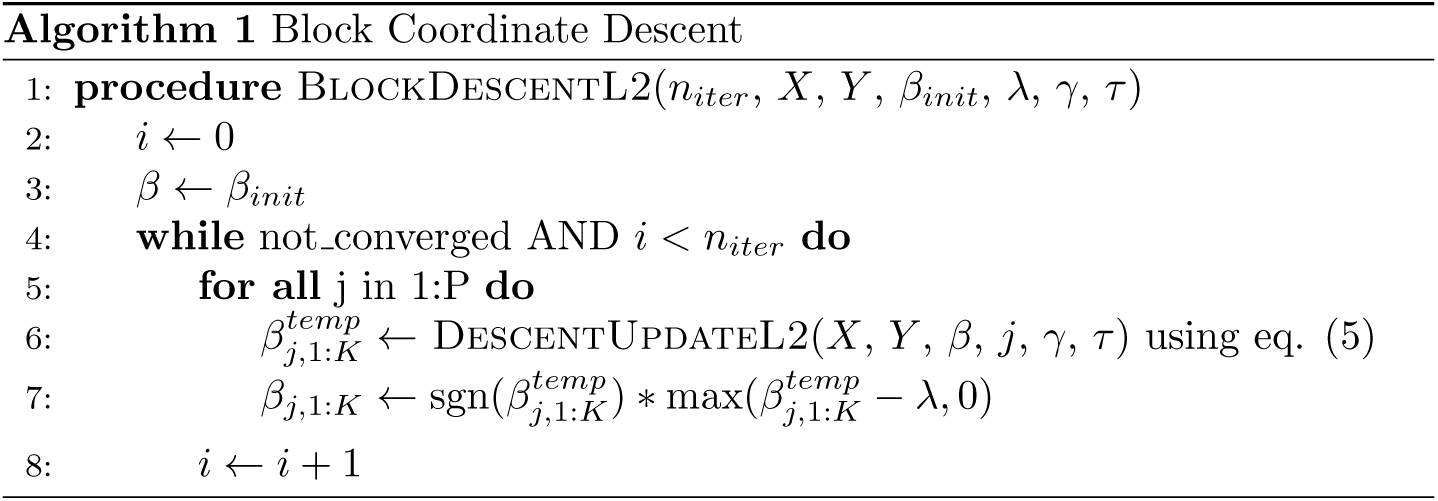

While Algorithm 1 is easy to understand and implement, a naive implementation in most programming languages will be still be slow due to the need for an inner for-loop over *p*, where *p* can be in the tens of thousands for the kinds of problems we will consider. In order to efficiently optimize *B*, we reformulate (1) as a classical lasso problem and apply the **glmnet** software [Friedman et al., 2010]. We transform the sum in first part of the objective into matrix form *y_flat_* — *X_diag_b_flat_* by defining *X_diag_* as a block-diagonal *n* × *pK* matrix with *X_k_* along the diagonals. The vector *b_flat_* is a flattened version of *B* with stacked *β_k_* vectors, and similarly for *y_flat_*. So we have:

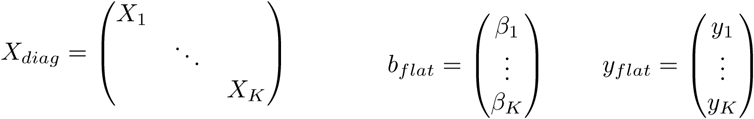

Now we move the *ℓ*_2_ fusion penalty into the first squared term by defining the augmented matrix 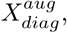 and augmented vector 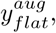 such that

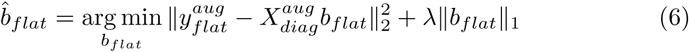

where

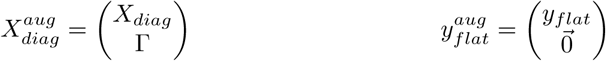

with Γ a *pK*(*K* — 1)/2 × *pk* matrix encoding the pair-wise fusion constraints, and 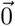 a *pK*(*K* — 1)/2 × 1 vector of zeros. Each block *Γ_k_*,_*k*_′, *k*, *k*′ *∈* [1, *K*], *k* < *k*′ of *p* rows of Γcorresponds to the fusion constraint between two coefficient vectors *β_k_* and *β_k_*′, with:

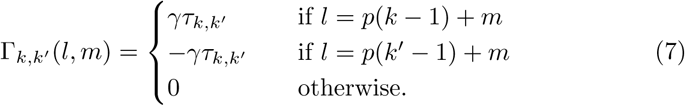

We can see that (6) is a classical lasso problem, to which `glmnet` can be directly applied.

#### 2.5.2 Proximal-Gradient Approach for Fused L1 Penalty

Optimising equation (2) by block gradient descent, while possible, is highly inefficient due to having to deal with the discontinuities in the objective function space. In Chen et al. [2010], the authors describe a proximal relaxation of this problem that introduces additional smoothing to turn the objective function *f_L_*_1_(B) into a continuously differentiable function 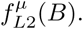. Chen et al. deal with the multi-task regression setting (with common *X* for each task); it is straightforward to adapt their procedure for the subgroup regression setting with different *X_k_* per subgroup.

It is notationally convenient to first introduce a graph formulation of the fusion penalties. We will think of the fusion constraints in terms of an undirected graph *G* = (*V*, *E*) with vertex set *V* = {1… *K*} corresponding to the subgroups and edges between all vertices. Then the *ℓ*_1_ penalised objective function can be written as:

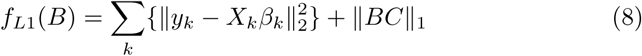

where the last term includes both sparsity and fusion penalties, via the matrix *C* = (*λI_k_*,*γH*), with *I_k_* the identity matrix of size *K*, *C* a *K* × |*E*| matrix 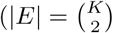 in this case) and *H_k_,_e_* = *τ*_*k*_,*Х* if if *e* = (*m*, Х) and *k* = *l*, *H_k_,_e_* = *—τ_m_,_k_* if *e* = (*m*, *Х*) and *k* = Х. Note that unlike in Chen et al. [2010], we require the explicit sum over *k* in the objective to account for different sample sizes *N_k_* in different groups^1^.

The graph formulation allows for zero edges by setting *τ_k_*,*_k_*′ to zero. We have implicitly assumed in the formulation of (1) and (2) that the relationship between subgroups is represented by an undirected graph. However, (8) is completely general, and it would be straightforward to incorporate a directed graph in our model. We have not pursued this avenue here, as there is no reason to suspect directionality in the subgroup relationships for the applications we consider below, and including directionality would double the number of tuning parameters *τ*_*k*_ *_k_*′ that need to be considered.

Following Chen et al. [2010], we can introduce an auxiliary matrix 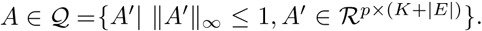. Because of duality between *ℓ*_1_ and *ℓ*_*∞*_, we can write 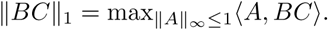. A smooth approximation of ∥*BC*∥_1_ is then obtained by writing:

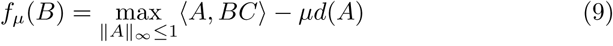

where *μ* is a positive smoothness parameter, and 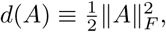 with ∥·∥*_F_* the Frobenius norm. They show that for a desired accuracy *∈*, we need to set 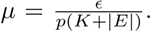. Theorem 1 in Chen et al. [2010] gives the gradient of *f_μ_*(*B*) as ∆*fμ*(*B*) = *A*C^T^*, where *A*\* is the optimal solution of (9). Replacing ∥*BC*∥_1_ by *f_μ_*(*B*) in equation (8), we obtain

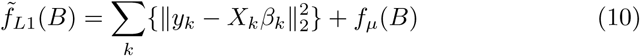

which is now continuously differentiable with gradient

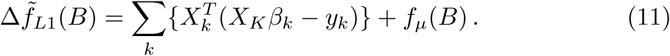

Chen et al. further show that *A*\* = *S*(*BC*/*μ*) where function S truncates each entry of *A*\* to the range [−1,1] to ensure that *A*\* *∈ Q*. An upper bound *L_U_* of the Lipschitz constant L can be derived as:

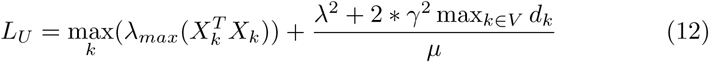

where *λ_max_(M)* is the largest eigenvalue of *M* and 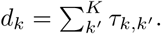.

With the derivation of the gradient in (11) and the Lipschitz bound in (12), we can now apply Nesterov’s method [Nesterov, 2005] for optimizing (10). The procedure is summarized in Algorithm 2. For more details on the proximal approach see Chen et al. [2010]

**Algorithm 2.**
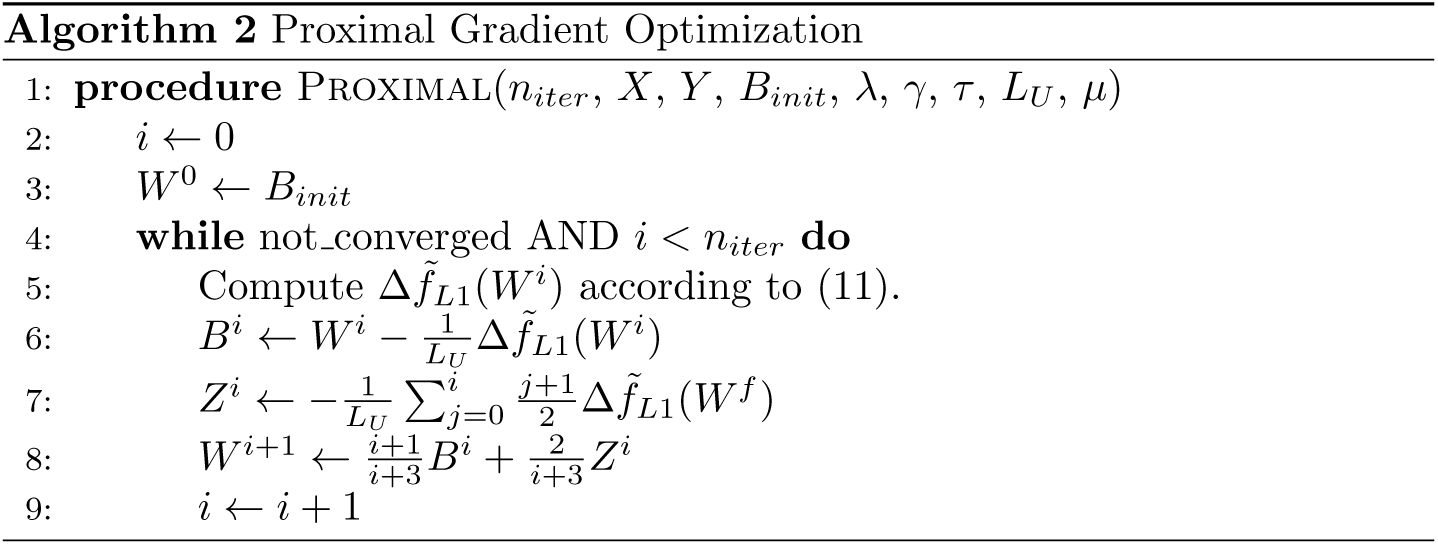

## 3 Simulation Study

To test the performance of the proposed approach, we simulated data from a model based on characteristics of a recent cancer dataset, the Cancer Cell Line Encyclopedia [CCLE; Barretina et al., 2012]. We treat cancer types as subgroups. To simulate data, we first estimated means and covariance matrices *μ_k_*,∑*_k_* for each of *K* = 9 subgroups (the eight cancer types with the latest sample sizes in CCLE plus a ninth for all other cancer types; covariances were estimated using the graphical lasso). For each group *k*, we then sampled covariates from the multivariate normal *Ν*(*μ_k_*,∑*_k_*).For a given total sample size *n*,subgroup sizes were consistent with those in the original data. We used a random subset of 200 gene expression levels (i.e. the dimensionality was fixed at *p* = 200). This parametric approach allowed us to vary sample sizes freely, including the case of total *n* larger than in the original dataset. The set-up is intended to roughly reflect the correlation structure of the covariates, but we do not expect it to capture all aspects of the real data.

We are interested in the situation in which it may be useful to share information between subgroups. But we are also interested in investigating performance in settings that do not agree with our model formulation (the extreme cases being where subgroups are either entirely dissimilar or identical). Let *V* = {1… *K*} be the set of subgroup indices (here, *K* = 9). We set regression coefficients to be identical in a subset *V*_0_ ⊆ *V* of the subgroups, such that the size *K*_0_ = |*V_0_*| of the subset governs the extent to which fusion could be useful. Specifically, if *K*_0_ = *K*, all subgroups have the same regression coefficients (i.e. favoring a pooled analysis using a single regression model) and at the other extreme if *K*_0_ = 1 all groups have differently drawn coefficients. Intermediate values of *K*_0_ give differing levels of similarity.

For a given value *K*_0_, we defined membership of *V*_0_ by considering the dif-ferences between the subgroup-specific models for the covariates. Specifically, we choose the *K*_0_ groups that minimized the sum of symmetrised KL divergences between subgroup-specific models. A coefficient vector was then drawn separately for each subgroup *k* ∉ *V*_0_ and one, shared coefficient vector drawn for all *k* ∈ *V*_0_. Each draw was done as follows. We first sampled a binary vector *b* of length *p* from a Bernoulli, i.e. *b_i_* ˜ Bernoulli(0.1). Then we drew *β_i_* ˜ *N_*trunc*_*(0,1) if *b_i_* = 1 and set *β_i_* = 0 otherwise, where *N_trunc_* (0,1) denotes a standard Normal with the interval (−0.1, 0.1) excluded (this is to ensure non-zero coefficients are not very small in magnitude). Note that in the case of *K*_0_ = 1, all groups have separately drawn coefficients and the between-subgroup KL divergence plays no role.

We compare our approaches with pooled and subgroup-wise analyses. These are performed using classical lasso (we use the **glmnet** implementation) on re-spectively the whole dataset or each subgroup separately.

Figure 1 shows performance when varying the number *k*_0_ of subgroups with shared coefficients, with the total number of samples fixed at *n* = 250. Here, a smaller value of *K*_0_ corresponds to less similarity between subgroup-specific coefficients in the underlying models. At intermediate values of *K*_0_ the fusion approaches offer gains over pooled and subgroup-wise analyses. This is because the pooled analyses are mis-specified due to the inhomogeneity of the data, while the subgroup-wise analyses, although correctly specified, must confront limited sample sizes since they analyze each subgroup entirely separately. In contrast, the fusion approaches are able to pool information across subgroups, but also allow for subgroup-specific coefficients. Importantly, even at the extremes of *K*_0_ = 1 (separately drawn coefficients for each subgroup) and *K*_0_ = 9 (all subgroups have exactly the same coefficients), the fusion approaches perform well. This demonstrates their flexibility in adapting the degree of fusion.

**Figure 1:**
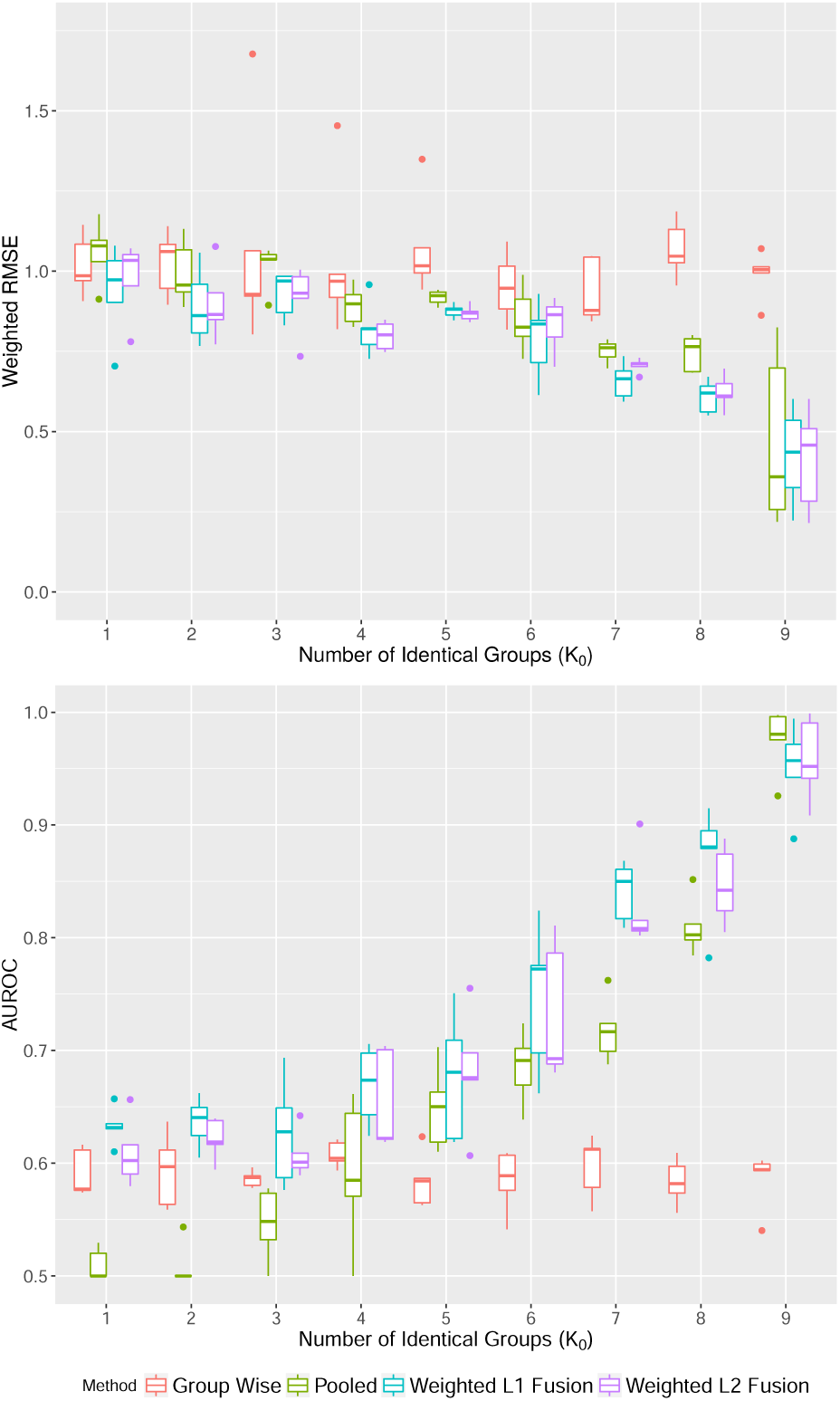
Simulated data, performance for varying *K*_0_. Here, the number of subgroups is fixed at *K* = 9 of which *K*_0_ have shared coefficients in the underlying data-generating model (see text for details of simulation set-up). A smaller *K0* corresponds to less similarity between underlying subgroup-specific models, with *K*_0_ = 1 representing the case where all subgroups have separately drawn coefficients while *K*_0_ = 9 represents an entirely homogenous model in which each subgroup has exactly the same regression coefficients. The total sample size is fixed at *n* = 250. Upper panel: Weighted root mean squared error (RMSE). RMSE is weighted by subgroup sizes. Lower panel: Area under the ROC curve (AUROC; with respect to the true sets of active variables with non-zero coefficients).

Figure 2 shows performance as a function of total sample size *n*. Here, the number of subgroups with identical coefficients is fixed at *K*_0_ = 4. This gives a relatively weak opportunity for information sharing, since 5/9 groups have separately drawn coefficients. Since the true *β*_k_’s are not identical, the pooled analysis is mis-specified and accordingly even at large sample sizes, it does not catch up with the other approaches. As expected, subgroup-wise analyses perform increasingly well at larger sample sizes. However, at smaller sample sizes the fusion approaches show some gains.

**Figure 2:**
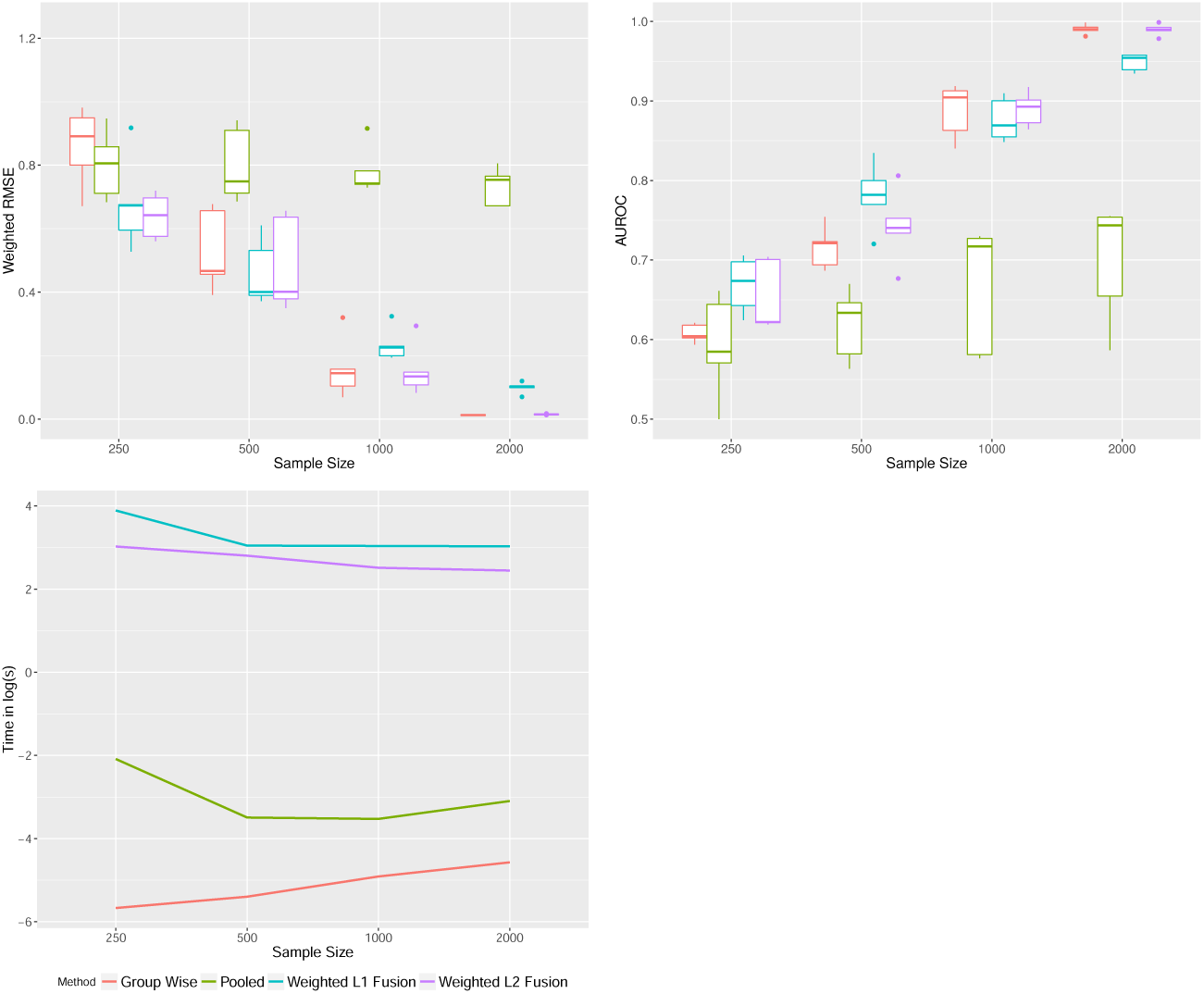
Simulated data, performance at varying sample sizes. Here, the number of subgroups is fixed at *K* = 9 of which *K*_0_ = 4 have shared coefficients in the underlying data-generating model (see text for details of simulation set-up). Top left: Root mean squared error (RMSE; weighted by subgroup sizes). Top right: Area under the ROC curve (AUROC; with respect to the true sets of active variables with non-zero coefficients). Bottom: Computational time taken in log seconds.

The *ℓ*_1_ and *ℓ*_2_ fusion approaches seem similar in performance. Our *ℓ*_2_ implementation leverages the `glmnet` package and is more computationally efficient than the*ℓ*_1_ approach. For computational convenience, in examples below we show results from the *ℓ*_2_ approach only.

## 4 Alzheimers disease: prediction of cognitive scores

Here, we use data from the Alzheimer’s Disease Neuroimaging Initiative (ADNI) [Mueller et al., 2005] to explore the ability of fusion approaches to estimate regression models linking clinical and genetic covariates to disease progression, as captured by cognitive test scores.

In 2014, ADNI made a subset of its data available for a DREAM challenge [Allen et al., 2016] and we use these data here. The dataset consists of a total of *n* = 767 individuals who were followed up over at least 24 months. Cognitive function was evaluated using the mini-mental state examination (MMSE). At baseline, individuals were classified as either cognitively normal (CN), early mild cognitive impairment (EMCI), late mild cognitive impairment (LMCI) or diagnosed with Alzheimer’s disease (AD). These form clinically-defined subgroups for our analysis. For the present analysis, we use only genetic data (single nucleotide polymorphisms or SNPs) and clinical profile as covariates and disregard the neuroimaging data.

The task is to predict the slope of MMSE scores over a 24-month period. The total number of SNPs available is ˜10^7^. Filtering by linkage disequilibrium reduces this to ˜2 × 10^6^. For computational ease, we pre-selected 20,000 of this latter group that gave the smallest residuals when regressed with the clinical variables against responses in the training set. We note that this biases our analyses, but we emphasize that our goal in this section is not biomarker discovery but comparison between approaches all using the same (pre-selected) covariates.

Figure 3 shows root mean squared error (RMSE) separately for each of the four subgroups. The fusion approaches offer substantial gains compared with pooled and subgroup-wise analyses (the latter performed very badly and are not shown in the figure). The biggest gain is for the AD subgroup. For the weighted fusion analysis the tuning parameters *τ_k_*, _*k*_′ were set using the distance between the means of each subgroup (in the space of genetic and clinical variables). Weighting did not appear to improve performance.

**Figure 3:**
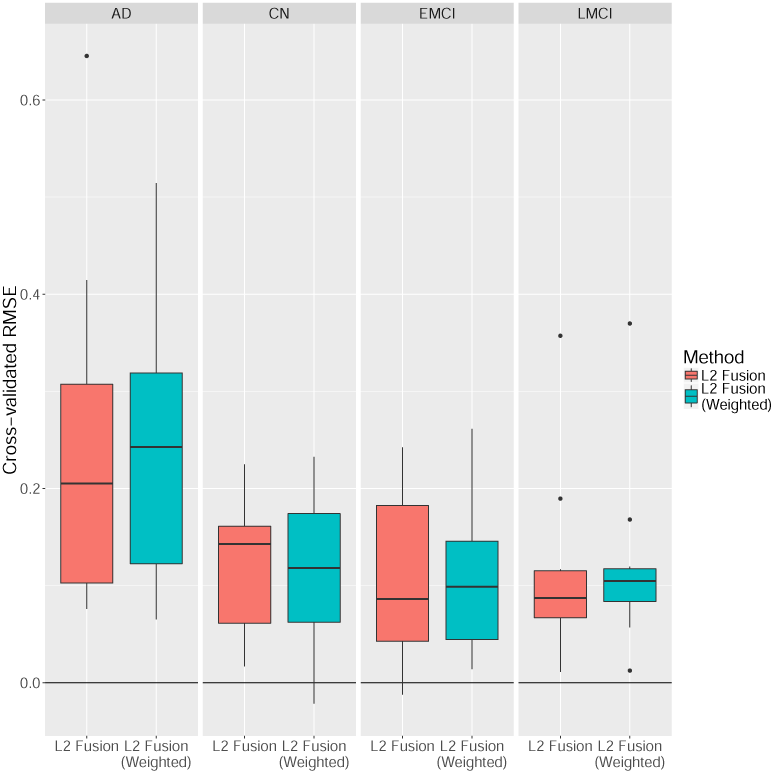
Alzheimers disease prediction results, ADNI data. Box plots showing difference in RMSE of fused methods compared to the pooled linear regression model (higher values indicate better performance by the fused methods). [Subgroup-wise analysis performed less well than pooled and is not shown; boxplots are over 10-fold cross-validation.]

Figure 4 shows scatter plots of predicted MMSE slopes versus the true slopes. The predictions shown were obtained in a held-out fashion via 10-fold crossvalidation (CV), as were the RMSE and Pearson correlations shown.

**Figure 4:**
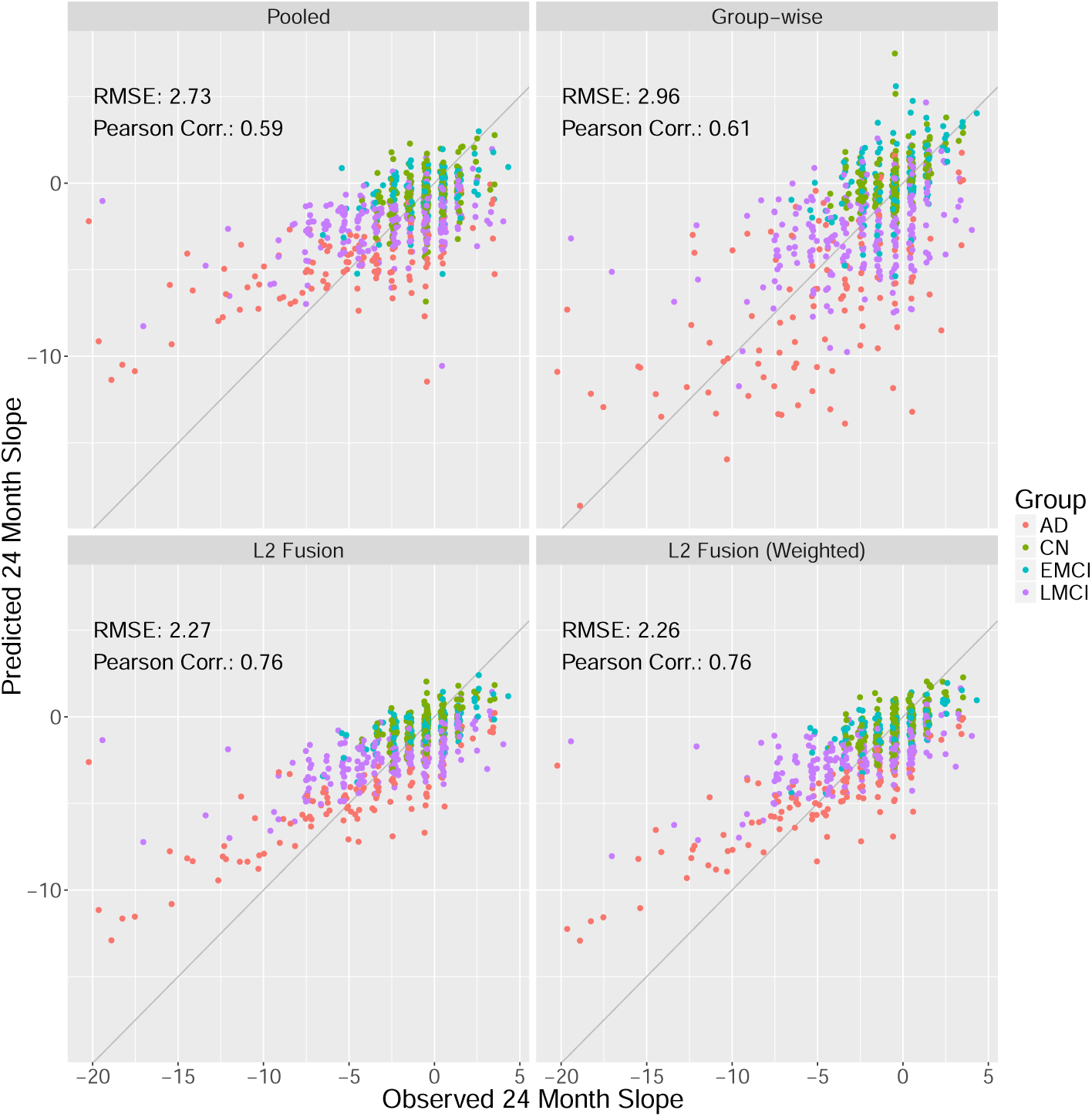
Alzheimers disease data, predicted vs. observed responses. Scatter plots show predicted and observed 24-month slopes for each of the standard and fused linear regression models. All predictions were obtained via 10-fold cross-validation.

Figure 5 shows a comparison of the estimated regression coefficients themselves. The subgroup-wise approach is much sparser than the other methods, likely due to the fact that it must operate entirely separately on each (relatively small-sample) subgroup. The pooled approach finds more influential variables but obviously there is no subgroup-specificity. The fused approach selects more variables than the subgroup-wise analysis, but there are many instances of subgroup-specificity in the estimates.

**Figure 5:**
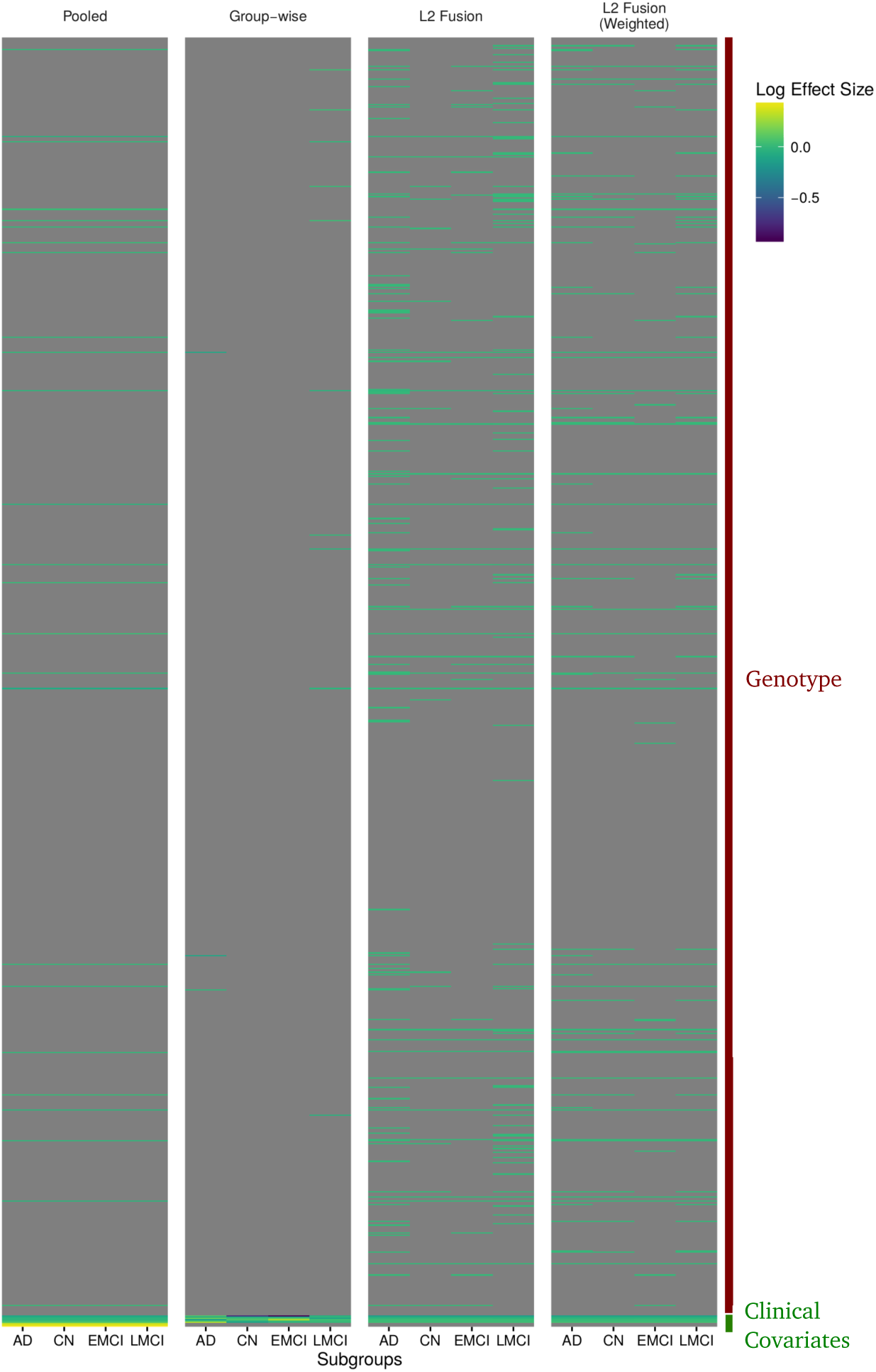
Alzheimers disease data, estimated regression coefficients. Heatmap showing estimated regression coefficients for the clinical variables and a representative subsample of the SNPs. Absolute coefficients are thresholded at *e*^−2^ to improve readability.

## 5 Prediction of therapeutic response in cancer cell lines

The Cancer Cell Line Encyclopedia [CCLE, Barretina et al., 2012] is a panel of 947 cancer cell lines with associated molecular measurements and responses to 24 anti-cancer agents. Here, we use these data to explore group-structured regression. We treat the area above the dose-response curve as the response and use expression levels of ˜20,000 human genes as covariates. We treat the cancer types as subgroups *k*. After discarding cell lines with missing values, we arrive at *n* ˜500 samples.

Figure 6 shows results over all 24 responses (anti-cancer agents). We observe that for most responses the L2 fusion approach shows either improved or similar prediction performance to pooled in terms of RMSE (weighted by subgroup size). Weighted fusion shows a similar performance to unweighted fusion.

**Figure 6:**
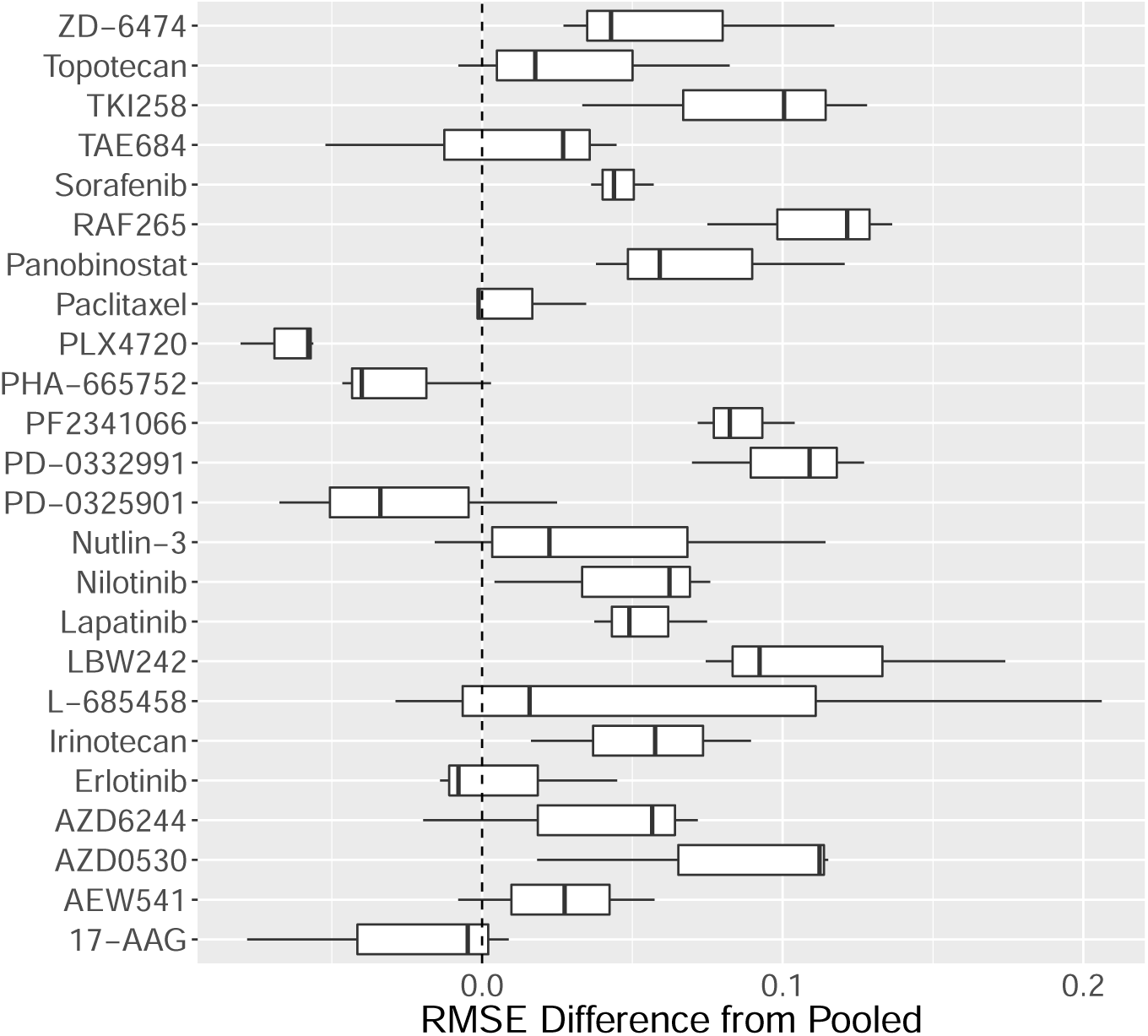
Cancer cell line therapeutic response prediction. Difference in weighted RMSE between L2 fusion approach and a pooled analysis. Results shown over 24 responses (anti-cancer agents) using data from the Cancer Cell Line Encyclopedia (CCLE); the dashed vertical line at zero indicates no difference, boxplots to the right indicate improvement (lower RMSE) over pooled.

Figure 7 shows results broken down by subgroup for two examples (responses PD-0332991 and PLX4720). In the former case, the fusion approaches largely outperform pooled and subgroup-wise analyses. In the second, pooled is the best performer, although the fusion approaches are similar in most subgroups.

**Figure 7:**
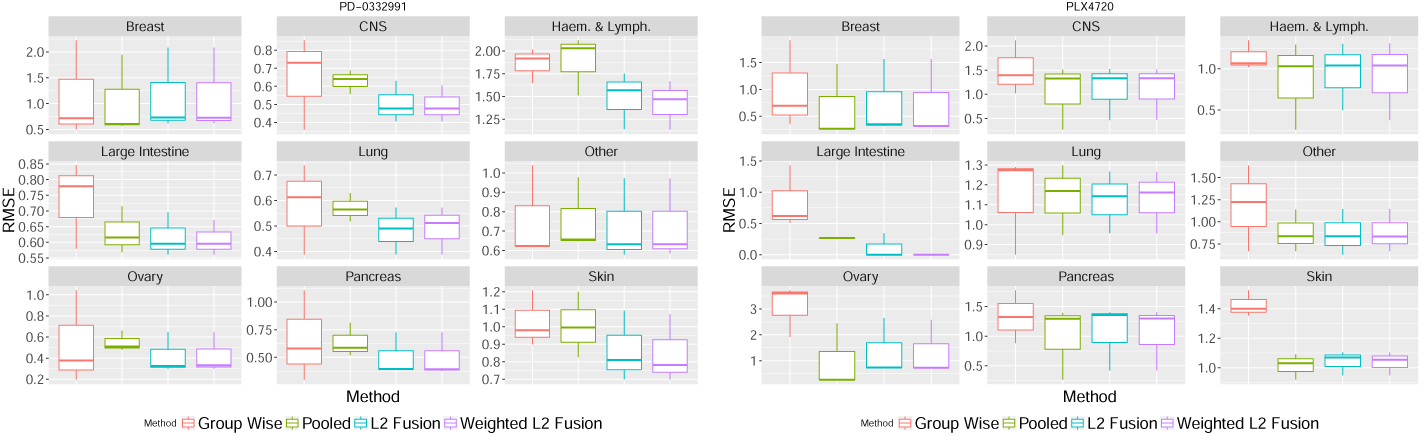
Cancer cell line therapeutic response prediction, broken down by subgroup (cancer type) for agents PD-0332991 and PLX4720.

## 6 ALS: prediction of disease progression

Amyotrophic lateral sclerosis (ALS) is an incurable neurodegenerative disease that can lead to death within three to four years of onset. However, about ten percent of patients survive more than 10 years. Prediction of disease progression remains an open question. We use data from the PROACT database, specifically data that were used in the 2015 DREAM ALS Stratification Prize4Life Challenge (data were retrieved from the PROACT database on 22/06/2015). Our aim is not to optimize predictive performance *per se* but rather to provide a case study exploring the use of fusion approaches in a moderate-dimensional, clinical data setting. In contrast to the Alzheimers example above, here the data are less high-dimensional and the subgrouping less clear cut (see below).

The data consist of observations from *n* = 2, 393 patients. Each patient was enrolled in a clinical trial and followed up for a minimum of 12 months after the start of the trial. Disease progression is captured by a clinical scale called the ALS Functional Rating Scale (ALSFRS). The task is to predict the slope of the ALSFRS score from 3 to 12 months (after the start of the trial). For each patient, available covariates include ALSFRS scores for the 0-3 month period, demographic information and longitudinal measurements of clinical variables. We follow the featurization and imputation procedures devised by Mackay [see Küffner et al., 2015] and obtain a total of *p* = 615 covariates.

Subgroups were defined as follows. The first subgroup consists of patients with disease onset before the start of the trial. The second subgroup consists of patients for whom onset was after the start of the trial and who have negative ALSFRS slope. The third subgroup of patients also had onset after the start of the trial but positive ALSFRS slope. Thus, the subgroups reflect severity of onset. As we believe that the pre-trial onset group are likely to differ most from the others, we manually set the distance between groups 1 and the other two groups to 1 (the maximum), and set the distance between groups 2 and 3 to 0.1.

Figure 8 shows (held-out) RMSEs by subgroup; we see that the largest improvement in prediction performance is in subgroup 1. The fusion approach leads to a modest improvement. The difference between weighted and unweighted fusion is negligible^2^.

**Figure 8:**
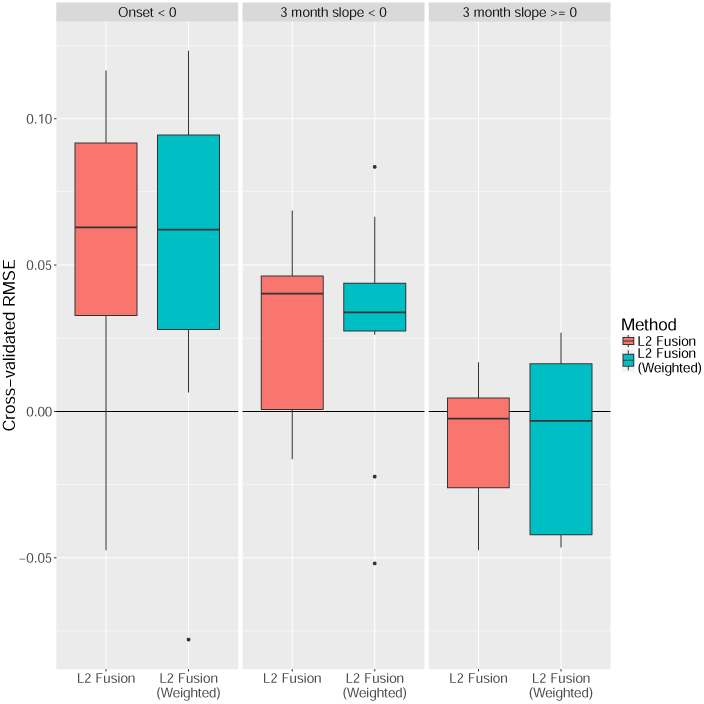
ALS prediction results. Box plots showing difference in RMSE of fused methods compared with the pooled linear regression model (higher values indicate better performance by the fused methods). [Subgroup-wise analysis performed less well than pooled and is not shown; boxplots are over 10-fold cross-validation.]

## 7 Conclusions

Many biomedical datasets are heterogenous, spanning multiple disease types (or other biological contexts) that are related but also expected to have specific underlying biology. This means that large datasets are often usefully thought of as comprising several smaller datasets, that have similarities but that cannot be assumed to be identically distributed. Statistically efficient regression in these group-structured settings requires ways to pool information where useful to do so, whilst retaining the possibility of subgroup-specific parameters and sparsity patterns. We proposed a penalized likelihood approach for high-dimensional regression in the group-structured setting that provides group-specific estimates with global sparsity and that allows for information sharing between groups.

In any given application, even when there are good scientific reasons to suspect differences in regression models between subgroups, it is hard to know in advance whether the nature of any differences is such that a specific kind of joint estimation would be beneficial. For example, if sample sizes are small and groups only slightly different, pooling may be more effective, or if the groups are entirely different, fusion of the kind we consider may not be useful. This means that in practice, either simple pooling or subgroup-wise analysis may be more effective than fusion. In our approach, the tuning parameter *γ* (set by crossvalidation) determines the extent of fusion in a data-adaptive manner, and we saw in several examples that this appears successful in giving results that are at worst close to the best of pooling and subgroup-wise analyses. For settings with widely divergent *n_k_*’s, it may be important to allow tuning parameters to depend on *n_k_* (we did not do so) and to consider alternative formulations that allow for asymmetric fusion.

An appealing feature of our approach is that it allows for subgroup-specific sparsity patterns and parameter estimates that may themselves be of scientific interest. We discussed point estimation, but did not discuss uncertainty quantification for these subgroup-specific estimates. A number of recent papers have discussed significance testing for lasso-type models [see e.g. Wasserman and Roeder, 2009, Lockhart et al., 2014, Städler and Mukherjee, 2016] and we think some of these ideas could be used with the models proposed here.

## 8 Software Availability

The R code used for the experiments in this paper has been made available as R package **fuser** on GitHub:https://github.com/FrankD/fuser. Scripts for reproducing the results in this paper can be obtained at:http://fhm-chicas-code.lancs.ac.uk/dondelin/SubgroupFusionPrediction.

## Footnotes

1 It would be possible to reformulate the first part of the objective in matrix form *Y_diag_* – *X_diag_ B_diag_* by defining *X_diag_* as a block-diagonal matrix as in Section 2.5.1, defining *B_diag_* as a *pK* ×*K* matrix with *β_k_* along the diagonals and similarly *y_diag_* as an *n* × *K* matrix with *y_k_* along the diagonals; however, this formulation is neither practical nor intuitive, and the gain in notational simplicity is negligible.

2 This dataset is different from the one reported in [Küffner et al., 2015], with larger variance in the slopes, and so RMSE values are not directly comparable; however, we note that performance for our methods compares favorably with that reported in the reference.

## 9 Acknowledgements

Data collection and sharing for the Alzheimer’s data application was funded by the Alzheimer’s Disease Neuroimaging Initiative (ADNI) (National Institutes of Health Grant U01 AG024904) and DOD ADNI (Department of Defense award number W81XWH-12-2-0012). ADNI is funded by the National Institute on Aging, the National Institute of Biomedical Imaging and Bioengineering, and through generous contributions from the following: AbbVie, Alzheimers Association; Alzheimers Drug Discovery Foundation; Araclon Biotech; BioClinica, Inc.; Biogen; Bristol-Myers Squibb Company;CereSpir, Inc.;Cogstate;Eisai Inc.; Elan Pharmaceuticals, Inc.; Eli Lilly and Company; EuroImmun; F. Hoffmann-La Roche Ltd and its affiliated company Genentech, Inc.; Fujirebio; GE Healthcare; IXICO Ltd.; Janssen Alzheimer Immunotherapy Research & Development, LLC.; Johnson & Johnson Pharmaceutical Research & Development LLC.; Lumosity; Lundbeck; Merck & Co., Inc.; Meso Scale Diagnostics, LLC.; NeuroRx Research; Neurotrack Technologies;Novartis Pharmaceuticals Corporation; Pfizer Inc.; Piramal Imaging;Servier; Takeda Pharmaceutical Company; and Transition Therapeutics. The Canadian Institutes of Health Research is providing funds to support ADNI clinical sites in Canada. Private sector contributions are facilitated by the Foundation for the National Institutes of Health (www.fnih.org). The grantee organization is the Northern California Institute for Research and Education, and the study is coordinated by the Alzheimers Therapeutic Research Institute at the University of Southern California. ADNI data are disseminated by the Laboratory for Neuro Imaging at the University of Southern California.

